# Inbreeding depression in polyploid species: a meta-analysis

**DOI:** 10.1101/2022.10.06.511129

**Authors:** Josselin Clo, Filip Kolář

**Author notes:** **For correspondence**: Josselin Clo, Faculty of Science, Charles University in Prague, Benátská 2, CZ-128 01 Prague, Czech Republic, ORCID 0000-0002-3295-9481. **Data**: Data and Rcode are available on GitHub (https://github.com/JosselinCLO/Meta_analysis_Inbreeding_depression).

## Abstract

Whole-genome duplication is a common mutation in eukaryotes with far-reaching phenotypic effects, the resulting morphological and fitness consequences and how they affect the survival of polyploid lineages are intensively studied. Another important factor may also determine the probability of establishment and success of polyploid lineages: inbreeding depression. Inbreeding depression is expected to play an important role in the establishment of neopolyploid lineages, their capacity to colonize new environments, and in the simultaneous evolution of ploidy and other life-history traits such as self-fertilization. Both theoretically and empirically, there is no consensus on the consequences of polyploidy on inbreeding depression. In this meta-analysis, we investigated the effect of polyploidy on the evolution of inbreeding depression, by performing a meta-analysis within angiosperm species. The main results of our study are that the consequences of polyploidy on inbreeding depression are complex and depend on the time since polyploidization. We found that young polyploid lineages have a much lower amount of inbreeding depression than their diploid relatives and their established counterparts. Natural polyploid lineages are intermediate, and have a higher amount of inbreeding depression than synthetic neopolyploids, and a smaller amount than diploids, suggesting that the negative effect of polyploidy on inbreeding depression decrease with time since polyploidization.

## Introduction

Polyploidization has occurred numerous times during the evolution of eukaryotes [1,2] and has been shown to have a broad range of phenotypic and genomic effects and to be an important mechanism for plant adaptation and speciation [3]. Nevertheless, polyploidization initially arises with several disadvantages in neopolyploids, like mitotic and meiotic dysfunction, genomic instability, decrease in fitness, and negative frequency-dependent selection [4–9].

Another important factor may also determine the probability of survival and success of polyploid lineages: inbreeding depression (ID hereafter) [10,11]. ID can be defined as the reduction of fitness found in selfed progenies compared to outcrossed progenies [12], and is predominantly due to the expression of recessive deleterious alleles at their homozygous state [13]. ID theoretically plays an important role in polyploids establishment. The initially low frequency of polyploid lineages within a diploid population may lead to strong bi-parental inbreeding [14]. In such conditions, it has been shown theoretically that a decrease in ID in polyploids compared to diploids is necessary for them to establish [10,11].

Polyploidy is theoretically expected to have both positive and negative effects on the amount of ID (see [15] for review). In the short term, it has been shown theoretically that in autopolyploid species, the strong bottleneck associated with polyploidization can strongly decrease the amount of ID due to the loss and/or the fixation of recessive deleterious mutations [16]. Even if not tested, we can expect similar consequences in allopolyploid species. In autopolyploid species with polysomic inheritance, even without a loss of genetic diversity, if frequencies of deleterious mutations remain similar in different cytotypes, as homozygosity increases at slower rates in autopolyploids compared to diploids [17], the expression of ID should be less severe in neoautopolyploids. In the long-term, the better masking of recessive deleterious mutations in autopolyploids should make them segregate at higher frequencies than in diploid populations [18]. Depending on the dominance coefficients of the mutations in the autopolyploid heterozygous genotypes, this increase in frequency can make ID to be smaller [19] or higher [20] in autopolyploids compared to diploids. Empirically, both a decrease [21,22] or an increase [23,24] of ID in synthetic and natural autopolyploids compared to diploids have been observed, and no strong consensus can be made. Allopolyploidy received much less attention, and there are consequently fewer expectations than in autopolyploid species. Hedrick [25] nevertheless showed that in homosporous, allopolyploid ferns, because offspring will be strongly homozygous due to intragametophytic selfing, ID should be lower in allopolyploids compared to diploids.

In this study, we investigated the effect of polyploidy on the evolution of ID. To do so, we performed a meta-analysis within angiosperm species. The main results of our study are that the effect of polyploidy on ID is complex and depends on the time since polyploidization. We found that synthetic polyploid lineages have a lower amount of ID than their diploid progenitors and their established counterpart. Natural polyploid lineages are intermediate, and have a higher amount of ID than synthetic neopolyploids, and a smaller amount than diploids.

## Material & Methods

### Dataset compilation

For this study, we were interested in the amount of ID in polyploids (auto- and allopolyploids, and of synthetic and natural genomic origins) compared to their diploid progenitors. As the studies comparing diploid and polyploid populations of the same species in the same article were rare, we extended the research to articles estimating the level of ID in polyploid populations, even without diploid controls.

We used Google Scholar, Web of Science, PubMed and Agricola databases in order to perform our literature survey. We used the keywords (“neopolyploid*” or “synthetic polyploid*” or “polyploid*”) and (“inbreeding”, “inbreeding depression”, “fitness”), and “plant*”. To be incorporated in the data collection, the selected study had to (1) define clearly the type (allo- or autopolyploidy) and level of ploidy of the population under study, (2) give the level of ID in the populations under study, or at least give the fitness of outbred and inbred progenies, such that we were able to infer ID ourselves, and (3) that ID equals *δ* = 1-(W_s_/W_o_) when W_s_ < W_o_ and *δ* = (W_o_/W_s_)−1 when W_s_ > W_o_, where W_s_ and W_o_ are respectively the fitness of selfed and outcrossed progenies [12]. Most of the time, the studies reported the numerical values in tables, but we sometimes had to extract the data directly from the figures, by using Plot Digitizer [26]. In the following part of the manuscript, we assumed that estimates found in synthetic polyploids (called neopolyploids in the following parts of the manuscript) will be our proxies for the short-term consequences of polyploidization on ID, while estimates from natural populations will be our proxies for the long-term consequences, even if the time since polyploidization is generally unknown and can greatly differ between species. A summary of the sampled species and articles can be found in Table 1.

**Table 1.** Summary of the sampled species, their ploidy levels, their genomic origin (auto- or allopolyploid), if they are natural or synthetical polyploids.

| Species | Ploidy levels | Genomic origin | Natural / synthetic | References |
| --- | --- | --- | --- | --- |
| *Acacia auriculiformis* | 2x, 4x | autopolyploid | synthetic | [41] |
| *Agropyron cristatum* | 2x, 4x | autopolyploid | natural & synthetic | [24,42] |
| *Amsinckia gloriosa* | 2x, 4x | autopolyploid | natural | [43] |
| *Anthericum liliago* | 2x, 4x | allopolyploid | natural | [44] |
| *Aster kantoensis* | 4x | autopolyploid | natural | [45] |
| *Beta vulgaris* | 4x | autopolyploid | natural | [46] |
| *Campanula americana* | 4x | autopolyploid | natural | [47,48] |
| *Centaurea stoebe* | 2x, 4x | allopolyploid | natural | [14] |
| *Chamerion angustifolium* | 2x, 4x | autopolyploid | natural & synthetic | [21,49–51] |
| *Clarkia davyi* | 2x, 4x | allopolyploid | natural | [52] |
| *Clarkia gracilis* | 2x, 4x | allopolyploid | natural | [52] |
| *Digitalis purpurea* | 4x | autopolyploid | natural | [53] |
| *Fragaria vesca* | 2x, 4x | autopolyploid | synthetic | [54] |
| *Iris versicolor* | 4x | allopolyploid | natural | [55] |
| *Jasione maritima* | 2x, 4x | autopolyploid | synthetic | [22] |
| *Knautia arvensis* | 4x | autopolyploid | natural | [56] |
| *Medicago sativa* | 4x | autopolyploid | natural | [23,57,58] |
| *Mercurialis annua* | 6x | autopolyploid | natural | [59,60] |
| *Pyrus communis* | 4x | autopolyploid | natural | [61] |
| *Scalesia affinis* | 4x | autopolyploid | natural | [62] |
| *Silene virginica* | 4x | autopolyploid | natural | [63] |
| *Spartina alterniflora* | 4x | allopolyploid | natural | [64] |
| *Trifolium hybridum* | 4x | autopolyploid | synthetic | [65] |
| *Vaccinium*<br>*corymbosum* | 2x, 4x | autopolyploid | natural | [66] |

### Considering phylogenetic non-independence

We tested if there was a potential phylogenetic correlation of our estimates in our analyses. To do so, we used the divergence times found in the TimeTree’s database [27], in order to reconstruct the phylogeny of our selected species. We then used the MEGA-X software [28] to transform the obtained matrix of distance into NEWICK format, with an UPGMA method. During this process, seven species were not found. We used the TimeTree database to find closely related species used for replacement of the missing ones (see Figure S1 and associated text to have the list of species). To test for a potential phylogenetic non-independence, we used the obtained phylogeny and ran linear mixed effect models by using the ‘metafor’ R package [29], and more precisely the rma.mv function that allows integrating phylogenetic correlation matrix into linear models. We tested if the amount of ID differed among diploid, neo- and natural polyploids using nested models. A first one in which a random-effect ‘Species’ is specified, and a second in which we included the phylogenetic correlation matrix as an additional random effect. The significance of the phylogenetic matrix was tested by performing a Likelihood Ratio Test (LRT) between the two abovementioned models.

### Inbreeding depression

We chose to perform a Bayesian meta-analysis, by using the MCMCglmm package [30]. We wrote the following model:

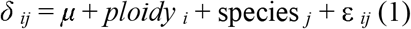

where *δ_ij_* is the level of ID, *μ* is the mean value, and *ploidy _i_* is the effect of the ploidy *i* (diploid, neo-polyploid [synthetic polyploid], and established tetraploid). As mentioned before, we only included a single random effect: species _*j*_ is the effect of the *jth* species, and ∊*_ij_* is the residual error. We assumed that the residual error followed a Gaussian distribution. We performed these models with two different datasets. The first one in which restricted to studies in which ID is estimated in diploid and/or neo- and natural polyploids simultaneously (called relatedness-controlled analysis after). In this dataset, we subtracted the ID level of diploids from the values of polyploids (Δ*δ* = *δ*_poly_ – *δ*_diplo_). If Δ*δ* < 0 (respectively > 0), it means that *δ* is smaller (respectively higher) in polyploids compared to diploids. In a second analysis, we included all studies and compared untransformed values of *δ* for diploids and polyploids (called complete analysis after). This analysis is potentially less robust, because we are comparing unrelated diploid and polyploid species, that can differ for other life-history traits potentially affecting ID levels.

For all analyses, we used the weakly informative, default priors proposed in MCMCglmm [30]. For fixed effects, the prior is a normal distribution with mean being equal to zero and a variance of 10^10^. For random effects, inverse-Wishart priors are implemented, with the degree of belief parameter being equal to zero and the expected variance being equal to 1. For all models, we used a burn period of 1.000.000 iterations, with a thinning interval of 50, and the MCMC chains were run for 6.000.000 iterations in total. The parameter models and associated 95% credible intervals were thus inferred from the sampling of the posterior distribution 100.000 times. We did a visual examination of the convergence, posterior traces and autocorrelation values of our models, as suggested in [30]. The trace of the sampled posterior, and posterior distributions for both models are available in supplementary materials (Figures S2 and S3).

## Results

The dataset was composed of 33 articles published between 1940 and 2020. These articles covered 25 species divided into 15 families of angiosperms. The relatedness-controlled analysis was based on 99 diploid-polyploids estimates of ID (70 in natural polyploid populations, 29 in synthetic polyploids). The complete analysis compiled 225 estimates of ID (195 in natural polyploid populations, 30 in synthetic polyploids). Most of the estimates have been estimated for autopolyploid species (187 in auto- and 23 in allopolyploid species). A summary can be found in Table 2.

**Table 2.** Summary of the number of estimates for the different categories of ploidy, for the relatedness-controlled and complete analyses.

| **Dataset** | **Diploids** | **Neo-polyploids** | **Natural polyploids** |
| --- | --- | --- | --- |
| **Relatedness-controlled** | 84 | 29 | 70 |
| **Complete** | 84 | 30 | 195 |

We found that the phylogenetic matrix did not improve the model (χ^2^ = 0.171, d.f. = 1, *p* = 0.680), so we decided to only keep the random effect ‘Species’. We found no differences between allo- and autopolyploid species (Tables S1, S2 & S3).

On average, polyploidy tended to decrease the amount of ID (Figure 1), with synthetic polyploids having the smallest mean ID level, and natural polyploids being intermediate between synthetic polyploids and diploids (Figure 1). In the relatedness-controlled analysis, synthetic polyploids had a significantly smaller level of ID than their diploid relatives (Figure 1A), but natural polyploids had an intermediate level, not significantly different from synthetic polyploids or diploids (Figure 1A). In the complete analysis, the amount of ID found in our restricted set of diploid estimates was in line with what was found in bigger studies (0.42 in [31], 0.38 [95% credible interval 0.27-0.60] in this study), confirming that we can use this value for comparisons. We found that all cytotypes had significantly different amounts of ID (Figure 1B). The synthetic polyploids had the lowest amount (Figure 1B, with a decrease of 69.0% compared to diploids), while natural polyploids were intermediate between the two other cytotypes (Figure 1B, with a decrease of 30.7% compared to diploids and an increase of 126.8% compared to synthetic polyploids).

**Figure 1.**
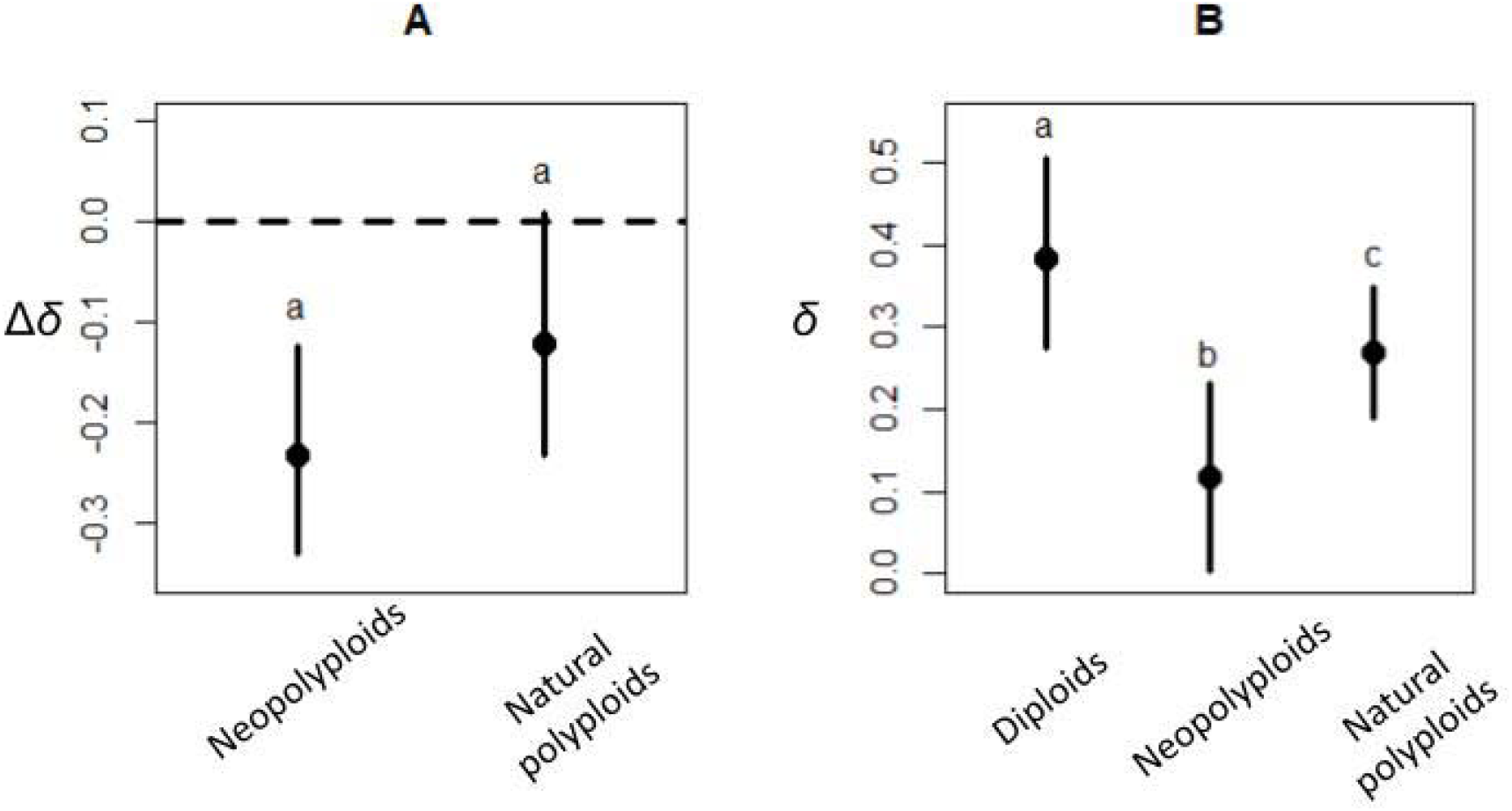
The evolution of inbreeding depression in polyploids (neo- and natural polyploids) compared to their diploid progenitors. **A**. Relatedness-controlled analysis, with Δ*δ* = *δ*_poly_ – *δ*_diplo_, a negative value showing that inbreeding depression is smaller in polyploids compared to diploids. Here, Δ*δ* is significantly different from zero for neopolyploids, but not for natural polyploids. **B**. Complete analysis. Different letters indicate significantly different mean values between ploidy levels. Error bars stand for the 95% credibility intervals.

## Discussion

### Inbreeding depression and establishment of new polyploid lineages

In this study, we found that ID decreases in polyploid populations compared to diploids. This result increases our understanding of how neopolyploid lineages can establish. Our meta-analysis confirmed theoretical expectations that due to an initial bottleneck [16], the masking of deleterious mutations that are in comparable frequencies as in diploids [18,32], and/or a slower increase in homozygosity during selfing events [17], new polyploid lineages benefit from a strong decrease in ID. Indeed, the small average amount of ID (*δ* = 0.119) found in synthetic polyploids suggests that potential bi-parental inbreeding should have a minor effect on the establishment probability of neopolyploids [10,11]. In the long-term, however, the amount of ID increases again in natural polyploids, as expected due to the increase in frequency of recessive deleterious mutations because of their better masking in polyploids compared to diploids[16,20]. Nevertheless, our results cannot conclude if natural polyploids have an intermediate or similar amount of ID to their diploid progenitors.

### The joint evolution of polyploidy, range expansion and mating system

Our results also give insights into why polyploidization can also lead to the evolution of other life-history traits. It has been observed that polyploid lineages can lead to the expansion of geographic [33] and climatic niches [34,35]. A theoretical argument is that a reduction in ID may favor such expansions, as biparental inbreeding, expected during the process due to bottleneck events [14], should have a minor effect in (neo)polyploids compared to diploids [11]. Our results support the theoretical prediction.

Finally, our results indicate that polyploidization could favor the transition from predominantly outcrossing to predominantly selfing mating systems. If an association between polyploidy and higher selfing rates has been found [36], it has been shown that this effect depends on the kind of ploidy. Husband *et al*. [21] showed that autopolyploid species generally self-fertilize less than diploid ones, while allopolyploids showed the opposite pattern [21,37]. Since our dataset is mainly composed of autopolyploid species (Table 1), our finding seems to be counterintuitive. Nevertheless, recent theoretical advances showed selfing only promotes autopolyploidization when neopolyploid lineages are at least as fit as their diploid counterparts [38], which is generally not the case [8,39].

Our results however suggest that geographic expansion and/or the evolution of selfing are more likely to occur in the very first generations following genome doubling, as ID is smaller in synthetic polyploids than in natural ones.

### Potential limitations

Even if informative, our study could suffer from potential bias. The first one is that synthetically produced polyploids could not be representative of natural young polyploid lineages. Nevertheless, the chemical treatments used the generate synthetic polyploids generally lead to a high to moderate death rate of treated diploids (see for example [40]), mimicking the expected bottleneck of genetic diversity that occurs during polyploidization events [16]. A second bias could be that the sampled diploid and natural polyploid lineages could differ in their selfing rate, which could be problematic if polyploids are more often predominantly selfers compared to diploids, as the observed decrease in ID could be due to an increased selfing rate that purges more efficiently deleterious mutations in natural polyploids compared to diploids [20]. Nevertheless, quick comparisons of the selfing rate found in our diploid and natural polyploid species suggest no differences (Table S4), and our dataset is mainly composed of autopolyploid species, that tend to have a smaller selfing rate than their diploid progenitors on average [21]. These results suggest that our analysis is conservative, and that the observed decrease in ID is due to the consequences of polyploidization *per se*. Finally, it is generally assumed that ID will affect the survival probability of polyploid populations, but such an association remains to be tested empirically.

## Conclusions

Our results are of primary importance for the understanding of polyploid establishment, and for the joint evolution of polyploidy with other life-history traits such as mating systems. However, our results remain preliminary, and further studies are needed to confirm the patterns described. Especially, theoretical and empirical studies in allopolyploid species would be of primary importance since there are few.

## Acknowledgement

We thank the associate editor and anonymous reviewers for helpful comments.

