## Supplementary materials for "Inbreeding depression in polyploid species: a meta-analysis"

**Table S1.** Average amount of inbreeding depression (and associated 95% credibility intervals) in diploids compared to neo- and natural auto- and allopolyploids.

| **Ploidy** | **Inbreeding depression** |
| --- | --- |
| Diploids | 0.356 [0.230 - 0.470] |
| Neo-autopolyploids | 0.114 [0.007 - 0.232] |
| Natural autopolyploids | 0.219 [0.028 - 0.380] |
| Natural allopolyploids | 0.320 [0.223 - 0.403] |

Table S2. Amount of inbreeding depression (and associated 95% credible intervals) for diploids, neopolyploids and natural polyploids in analyses including or excluding allopolyploid species, in the relatedness-controlled analysis.

| **Ploidy** | **Including allopolyploids** | **Excluding allopolyploids** |
| --- | --- | --- |
| Δδ **Neopolyploids** | -0.231 [-0.329 - -0.124] | -0.220 [-0.338 – -0.118] |
| Δδ **Natural polyploids** | -0.121 [-0.232 - 0.010] | -0.103 [-0.228 - 0.040] |

Table S3. Amount of inbreeding depression (and associated 95% credible intervals) for diploids, neopolyploids and natural polyploids in analyses including or excluding allopolyploid species, in the complete analysis.

| **Ploidy** | **Including allopolyploids** | **Excluding allopolyploids** |
| --- | --- | --- |
| **Diploids** | 0.384 [0.274 - 0.596] | 0.384 [0.274 - 0.596] |
| **Neopolyploids** | 0.119 [0.003 - 0.232] | 0.140 [0.024 - 0.263] |
| **Natural polyploids** | 0.270 [0.190 - 0.350] | 0.300 [0.203 - 0.390] |

**Table S4.** Average selfing rate found in diploid and polyploids species.

| **Ploidy** | **Selfing rate** |
| --- | --- |
| Diploids | 0.195 |
| Polyploids | 0.163 |
| *t*-test: *t = 1.191, d.f. = 140, p-value = 0.236* | |

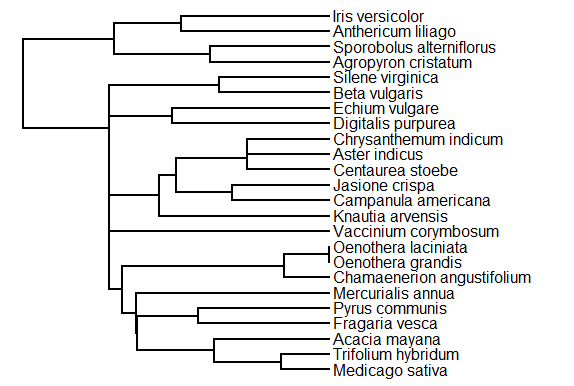

**Figure S1.** Phylogeny of the species used in the meta-analysis. We reconstructed the phylogeny of all sampled species, using the divergence times from the TimeTree’s database [1]. The obtained distance matrix was then transformed into NEWICK format using the UPGMA method implemented in the MEGA-X software [2]. During this process, seven species were not found. We replaced them by closely related species from the same genus found in the TimeTree database. The changes made are *Jasione maritima* by *J. Crispa*, *Amsinckia gloriosa* by *Echium vulgare*, *Scalesia affinis* by *Chrysanthemum indicum*, *Clarkia davyi* by *Oenothera grandis*, *C. gracilis* by *O. laciniata*, *Aster kantoensis* by *A. indicus*, *Acacia auriculiformis* by *A. mayana*.

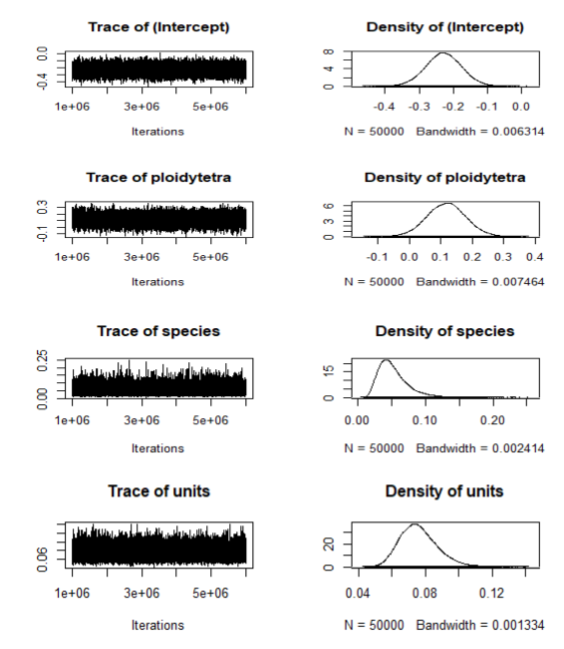

**Figure S2.** Summary plot of the Markov Chain for fixed and random effects of the relatedness-controlled analysis. The left plots are traces of the sampled posteriors. The right plots are the posterior distributions.

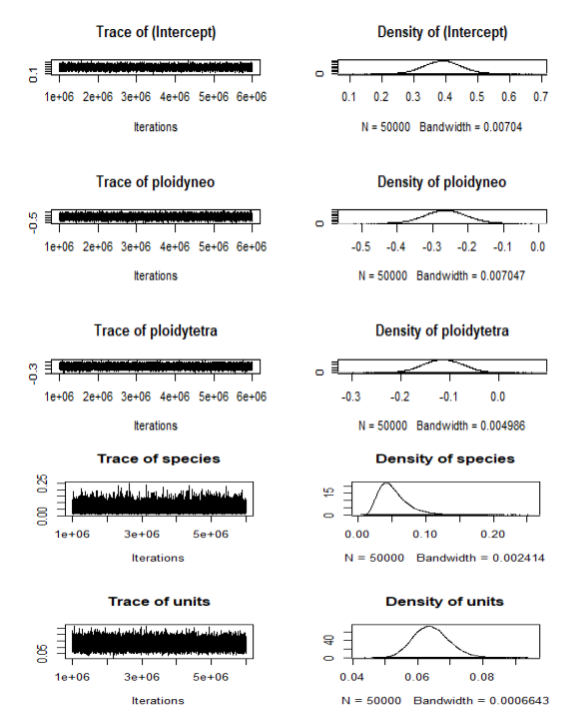

**Figure S3.** Summary plot of the Markov Chain for fixed and random effects of the complete analysis. The left plots are traces of the sampled posteriors. The right plots are the posterior distributions.
